# Protein language models are performant in structure-free virtual screening

**DOI:** 10.1101/2024.04.16.589765

**Authors:** Hilbert Yuen In Lam, Jia Sheng Guan, Xing Er Ong, Robbe Pincket, Yuguang Mu

## Abstract

Hitherto virtual screening has been typically performed using a structure-based drug design paradigm. Such methods typically require the use of molecular docking on high-resolution three-dimensional structures of a target protein - a computationally-intensive and time-consuming exercise. This work demonstrates that by employing protein language models and molecular graphs as inputs to a novel graph-to-transformer cross-attention mechanism, a screening power comparable to state-of-the-art structure-based models can be achieved. The implications thereof include highly expedited virtual screening due to the greatly reduced compute required to run this model, and the ability to perform early stages of computer-aided drug design in the complete absence of 3D protein structure.

## 2.1 Introduction

One major pillar of rational drug design has always been the use of virtual screening through docking in what is commonly known as structure-based drug design (SBDD) [1]. In docking, molecular ligands are typically conformationally explored in a protein pocket either through the use of biophysically defined constraints or machine learning (ML) methods, and a best pose with its corresponding computed binding affinity reported. A quintessential virtual screening pipeline will iteratively perform docking through a library, usually consisting of millions to billions of unique chemical compounds, and rank the ligands based on the derived affinity - the top scored ligands will then proceed onto the next phase of drug development, either through computational means such as molecular dynamics (MD) simulations or through experimental validation [2].

In order to enhance the accuracy of SBDD and virtual screening (VS), significant progress has been made in developing docking tools and rescoring functions - the latter of which are typically deep learning methods that, given a docked ligand, output a correction term or a new score entirely. These re-scoring methods have shown large promise, increasing the screening power in benchmarks [3–5]. Screening power is defined as the ability of a given model to differentiate between what will and will not bind to a target experimentally [6], and is largely considered the ultimate test of any virtual screening model. By increasing screening power, a model can more effectively discover leads, and overall improve the efficacy of computer-aided drug design (CADD).

However, docking itself and the addition of a rescoring term greatly increases the computational time and expense required to score each ligand. With very large libraries being used and an estimated chemical space of 10^63^ unique compounds [7], this increased compute in already time-consuming VS means that either cost would have to go up per target or the amount of ligands that can be screened per target would have to go down, inadvertently compromising the comprehensiveness of a virtual screening pipeline on the chemical space explored.

Aggravatingly, recent work also suggests that many SBDD deep learning models merely memorise the ligands, resulting in poor generalisability and outcomes [8]. In the case of models predicting the binding affinity directly, studies have shown that many of these models consider less of the protein-ligand interaction but more so of quantitative structure analysis relationship (QSAR) of the ligand alone in making their predictions [9]. Intrinsic biases have also been found in common datasets used for training SBDD deep learning models [9]. There is also the issue of limited data, with common datasets such as the PDBBind+ dataset [10] consisting of 22,920 ligand-protein pairs, further complicating the issue of training performative and generalisable models due to data scarcity. SBDD also has a pitfall in which only a single snapshot of a pocket is considered during docking and subsequent rescoring - and although these limitations have been addressed through many works throughout the years [11], the fundamental flexible and dynamic nature of the protein is still, unfortunately, largely given little consideration [12]. The performance of molecular docking can also vary between different conformational states of a protein (i.e. the apo or holo state), and hence docking predominantly suffers from considering the induced fitting of ligands [13].

To address this, some have resorted to instead predicting drug-target affinity (DTA), an experimentally derived figure such as the half-maximal inhibitory concentration (IC_50_) or dissociation constants. Such work includes SSM-DTA [14], a sequence-based deep learning method that achieves state-of-the-art performance in DTA prediction. However, in this work, the authors demonstrate that DTA-only models do not necessarily have the highest screening power.

Therefore, to tackle the issue of limited data in SBDD, the single snapshot issue, the lack of consideration of the entire protein during SBDD, induced fitting model and different conformations, this work combines protein language models (PLMs), a form of large language model (LLM) together with graph neural networks to represent ligands to perform virtual screening without inputting structural information (Fig. 1). The fundamental idea is that any structural and dynamic information of proteins is implicit in any PLMs pre-trained through self-supervised means such as Evolutionary Scale Modelling 2 (ESM-2) and this theorem has been further demonstrated in works such as ESMFold [15]. This work further introduces a novel graph-to-transformer cross-attention block, which essentially treats every single node in a graph as a token and allows it to query a separate sequence. The combined model, named **B**inding **IN**teraction **D**etermination (BIND), achieves comparable performance to state-of-the-art SBDD models with a fraction of the compute and time required and with only protein sequence and ligand information as inputs. Primarily, this work differs from previous work in its use of a decoy/true binder classifier and application to datasets typically used to benchmark SBDD models such as DUD-AD [16]. Through the use of this model, a state-of-the-art reverse screening performance was also achieved when compared with SBDD models on AlphaFold2-predicted structures. This work also showed that when trained with the multi-objective classification and DTA as used in BIND, the model’s DTA prediction also outperforms state-of-the-art DTA-only models in the screening power domain. Lastly, the trained model in this work can be deconstructed to determine which residues of a protein contribute most to the protein-ligand interaction, despite never being trained to do so.

**Figure 1.**
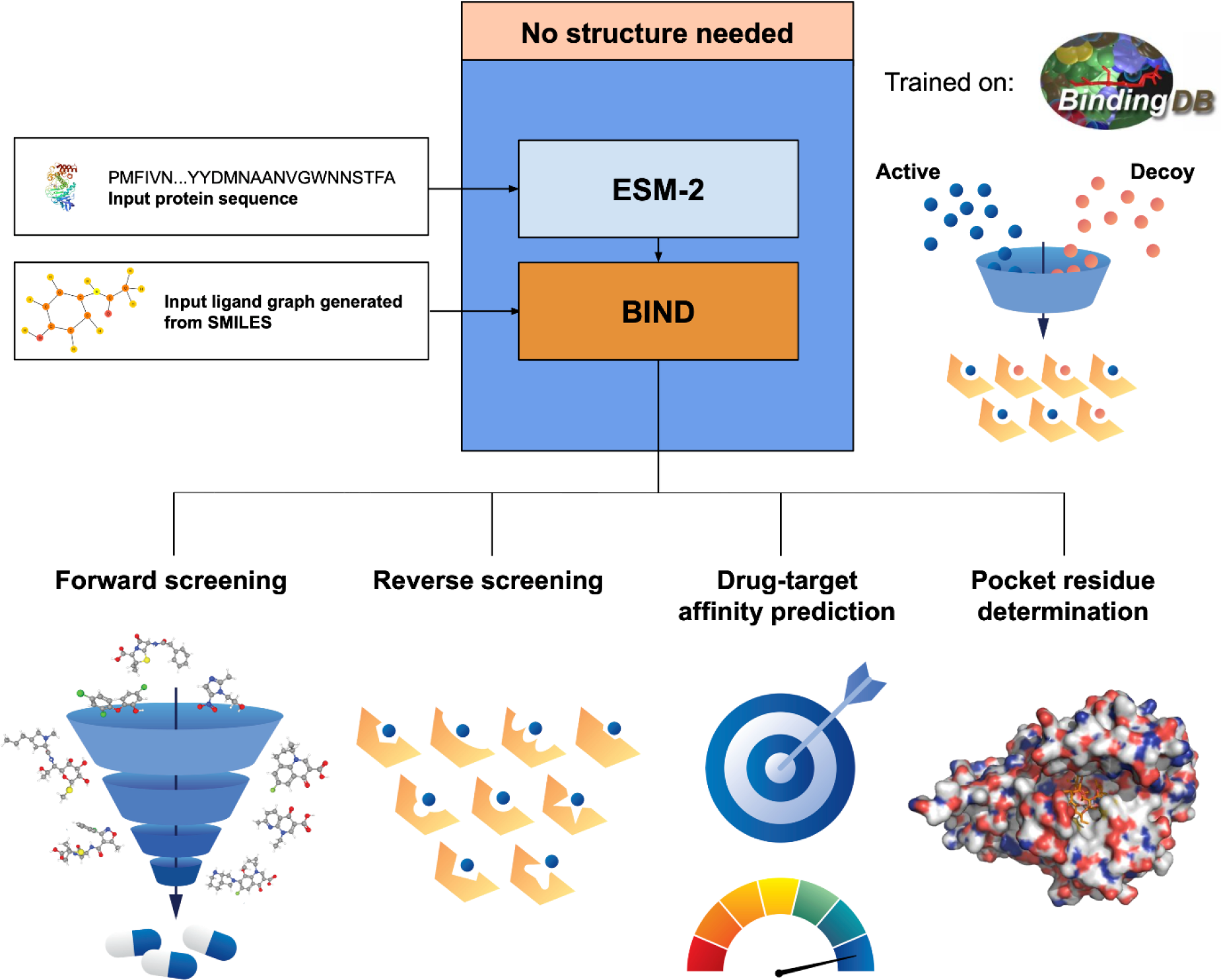
BIND is able to perform forward screening, reverse screening, drug-target affinity prediction, and elucidate residues that make up the pocket, all without structural input. BIND is trained on the BindingDB dataset only consisting of protein sequences and experimentally determined DTA values. By attaching BIND to a pre-trained ESM-2 protein language model and feeding in a protein sequence and SMILES molecular representation, which are then converted into graphs, the model can effectively discriminate between active and decoy ligands. This allows BIND to be used in forward and reverse screening, whilst predicting DTA values. Furthermore, residues in the binding pocket can be roughly elucidated using BIND, despite the model never being trained to do so.

## 3. Results

### 3.1 Screening power of BIND comparable to top models

The screening power of the model (Fig. 2) is similar to that of top SBDD models for all benchmarks performed (Table 1-5). Overall, a high enrichment factor is observed for all the benchmarks tested. In the DEKOIS 2.0, DUD-AD, DUD-E and LIT-PCBA datasets, the BIND model appears to have the highest enrichment factor at 1% cutoff with respect to reviewed literature (Fig. 3c-f). However, the model falls short in the CASF-2016 forward screening dataset (Fig. 3a-b), achieving higher enrichment than Glide-SP and Glide-XP but is overall deficient in the success rate and when compared to other very recent machine learning methods. Furthermore, the model’s enrichment factor is higher across all datasets tested when compared to using the pIC_50_ prediction from SSM-DTA, a fully DTA prediction model (Fig. 1) . The training and validation losses of the evaluated model are described in Supplementary Fig. 1, and individual benchmark scores for each protein are given in Supplementary File 1. Further training experiments to optimise the model and their corresponding results are also available in the supplementary material. In zero-shot protein settings, where the sequences in the training set containing > 90% homology compared to the evaluation datasets and their associated ligands are removed, the zero-shot BIND model still achieves good performance, outperforming some contemporary score functions in the datasets tested. The individual zero-shot BIND results are given in Supplementary File 2.

**Figure 2.**
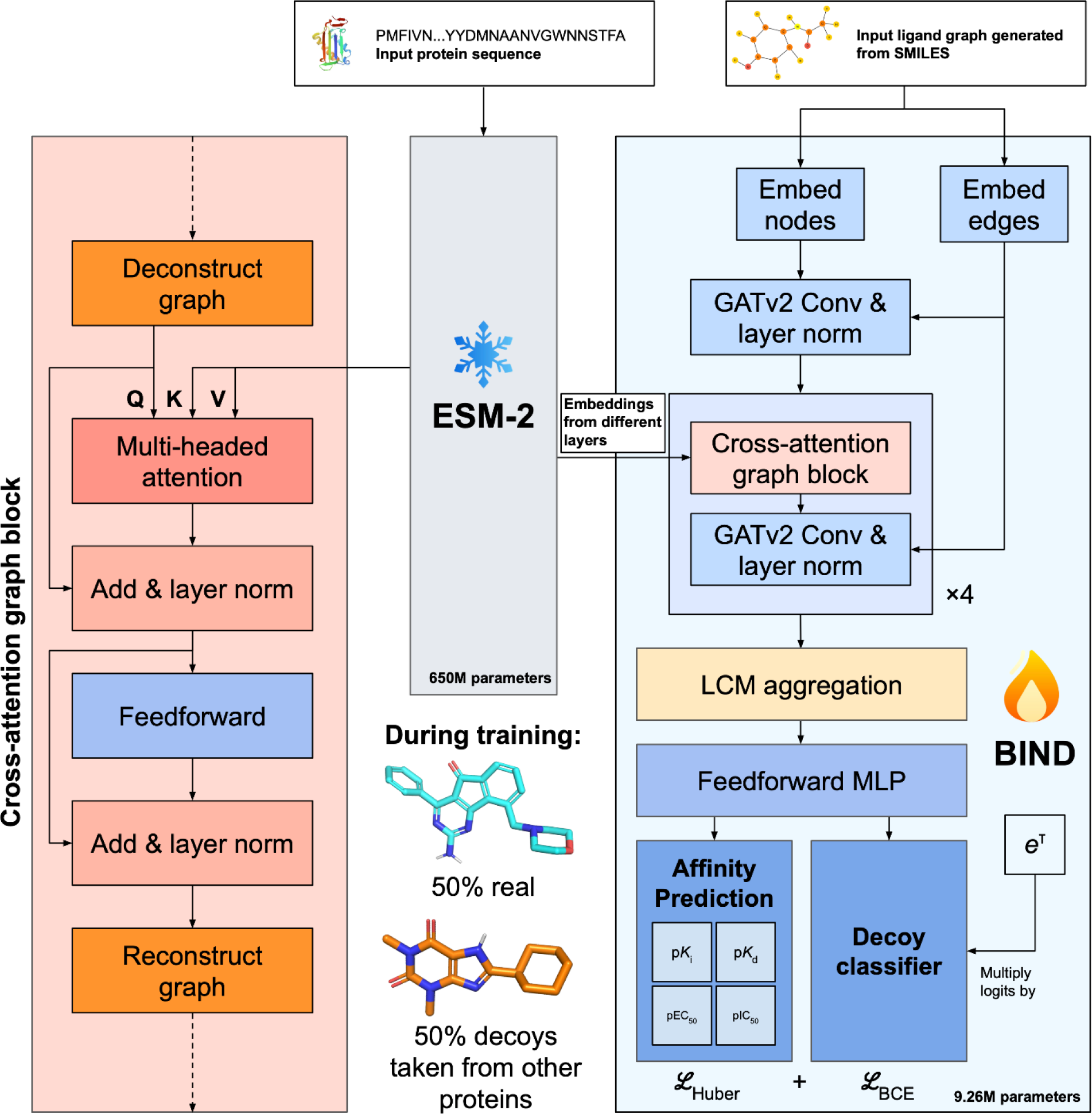
BIND architecture incorporates a proposed cross-attention graph block, and is trained with both true ligands and decoys taken from other proteins in the same dataset. The cross-attention graph block essentially deconstructs a graph and treats each node as a token for cross-attention - this allows the ligand to “query” the protein and its important parts. The loss is a summation of Huber loss for the affinity predictions, and binary cross entropy loss for the decoy classifier. During training, ESM-2’s weights are frozen such that only the 9.26M parameters from BIND are tuned. Q, K, and V in the diagram represent the query, key and value inputs characteristic of transformers.

**Figure 3.**
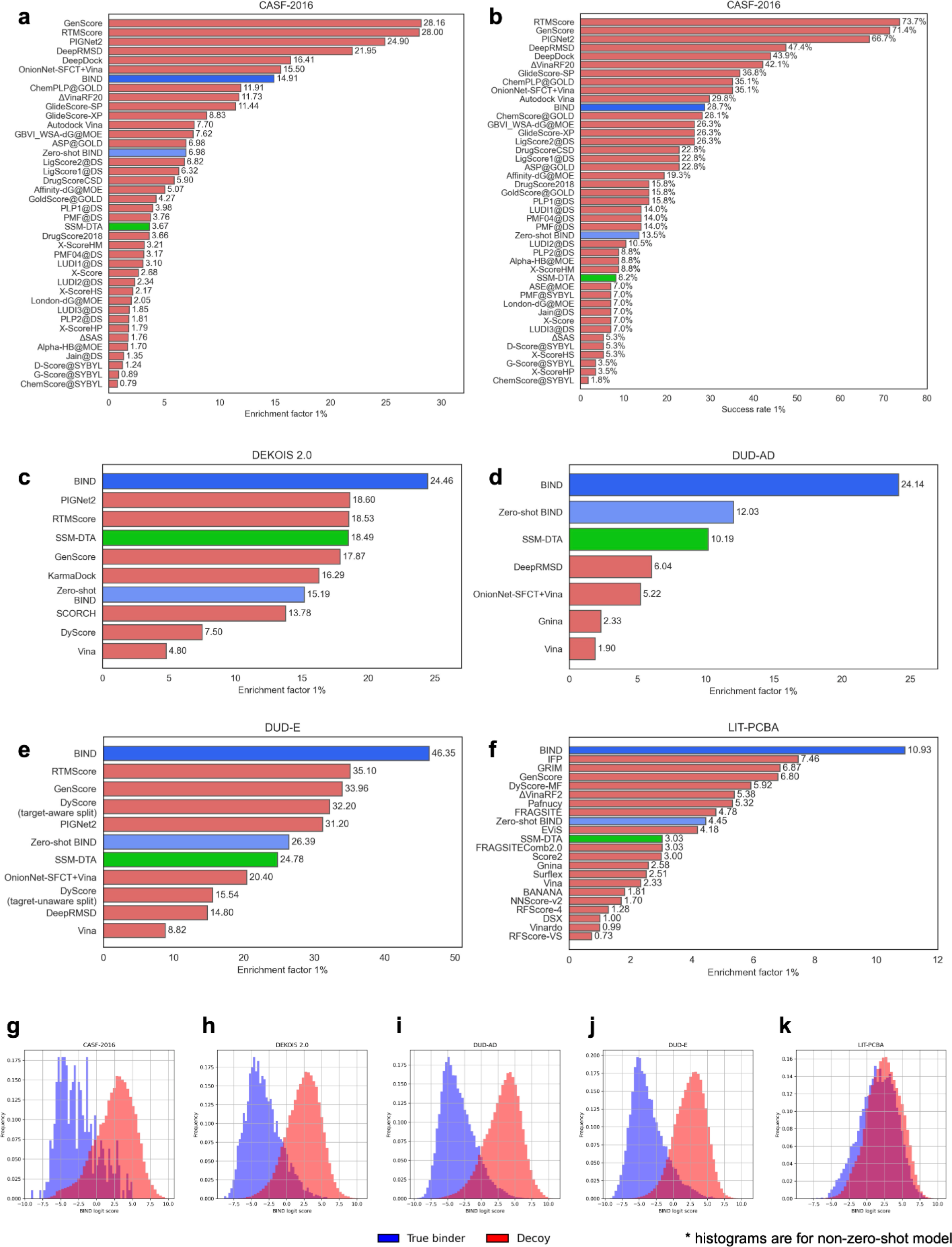
The model is performant on docking benchmarks and outperforms the state-of-the-art DTA only model in enrichment. CASF-2016, DEKOIS 2.0, DUD-AD, DUD-E and LIT-PCBA were evaluated. (a-b) CASF-2016 1% enrichment factors and 1% success rate, (c-f) DEKOIS 2.0, DUD-AD, DUD-E and LIT-PCBA 1% enrichment factors, (g-k) probability-normalised histogram of the non-zero-shot BIND classifier logit output showing separation between true binders and decoys. All enrichment factors and success rates are averaged across the entire datasets evaluated, and histograms shown are distributions of logit scores for all proteins and ligands tested. The green bar indicates the enrichment factor of the published SSM-DTA model, which predicts the pIC_50_ DTA. The zero-shot BIND model indicates the model in which proteins with > 90% homology comparative to the evaluation datasets are removed during training.

**Table 1.** Screening power results for top models and commonly used models on CASF-2016, with all scores represented as the mean. Arrows indicate direction of better results.

| Score function / model | Method | Prediction target | EF<sub>1%</sub> ↑ | Forward screening success rate 1% ↑ |
| --- | --- | --- | --- | --- |
| GenScore [42] | Deep learning, structure-based | Distance likelihood | 28.20 | 71.9% |
| RTMScore [5] | Deep learning, structure-based | Distance likelihood | 28.00 | 66.7% |
| PIGNet2 [43] | Deep learning, physics-based, structure-based | Binding affinity | 24.90 | 66.7% |
| DeepRMSD [44] | Deep learning, structure-based | $\Delta$ binding affinity | 21.95 | 47.4% |
| DeepDock [45] | Deep learning, structure-based | Distance likelihood | 16.41 | 43.9% |
| OnionNet-SFCT [4] | Deep learning, structure-based | $\Delta$ binding affinity | 15.5 | 35.1% |
| BIND | Deep learning, sequence-based | Binary classification between true binders and decoys | 14.91 | 28.7% |
| ChemPLP@GOLD [46] | Physics-based, Structure-based | Binding affinity | 11.91 [6] | 35.1% [6] |
| ΔVinaRF<sub>20</sub> [47] | Machine learning, structure-based | $\Delta$ binding affinity | 11.73 [6] | 42.1% [6] |
| Glide-SP [48] | Physics-based, structure-based | Binding affinity | 11.44 [6] | 36.8% [6] |
| Glide-XP [48] | Physics-based, structure-based | Binding affinity | 8.83 [6] | 26.3% [6] |
| ChemScore@GOLD [49] | Machine learning, structure-based | Binding affinity | 8.65 [6] | 28.1% [6] |
| Autodock Vina [50] | Physics-based, structure-based | Binding affinity | 7.70 [6] | 29.8% [6] |
| Zero-shot BIND (90% homologous sequences removed) | Deep learning, sequence-based | Binary classification between true binders and decoys | 6.98 | 13.5% |
| SSM-DTA (pIC<sub>50</sub>) [14] | Machine learning, sequence-based | Drug-target affinity | 3.67 [6] | 8.2% [6] |

\* best model's results as reported by respective authors

**Table 2.**
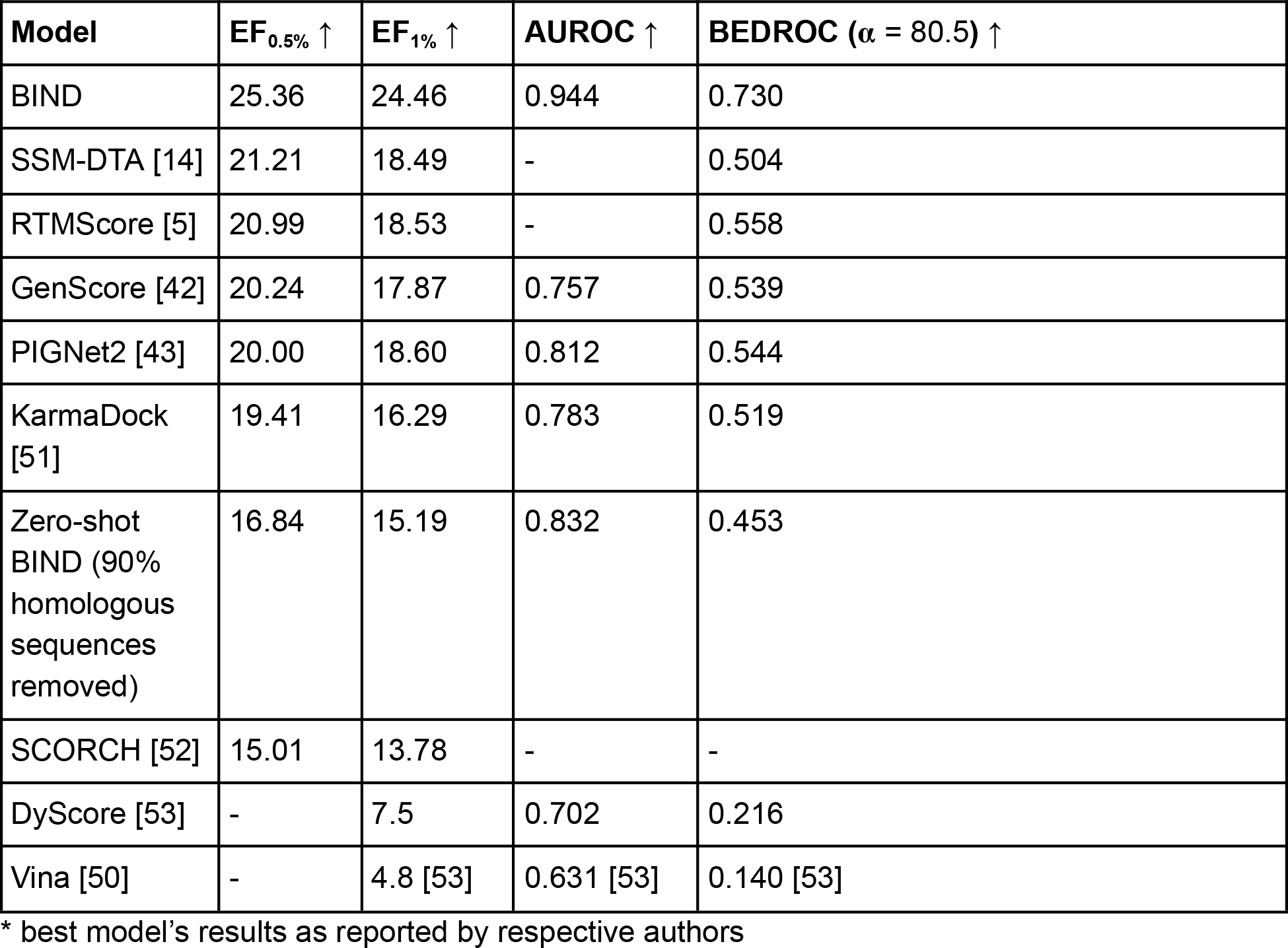
Screening power results for DEKOIS 2.0, with all scores represented as the mean. Arrows indicate direction of better results.

| Model | EF<sub>0.5%</sub> ↑ | EF<sub>1%</sub> ↑ | AUROC ↑ | BEDROC ($\alpha = 80.5$) ↑ |
| --- | --- | --- | --- | --- |
| BIND | 25.36 | 24.46 | 0.944 | 0.730 |
| SSM-DTA [14] | 21.21 | 18.49 | - | 0.504 |
| RTMScore [5] | 20.99 | 18.53 | - | 0.558 |
| GenScore [42] | 20.24 | 17.87 | 0.757 | 0.539 |
| PIGNet2 [43] | 20.00 | 18.60 | 0.812 | 0.544 |
| KarmaDock [51] | 19.41 | 16.29 | 0.783 | 0.519 |
| Zero-shot BIND (90% homologous sequences removed) | 16.84 | 15.19 | 0.832 | 0.453 |
| SCORCH [52] | 15.01 | 13.78 | - | - |
| DyScore [53] | - | 7.5 | 0.702 | 0.216 |
| Vina [50] | - | 4.8 [53] | 0.631 [53] | 0.140 [53] |

\* best model's results as reported by respective authors

**Table 3.** Screening power results for DUD-AD, with all scores represented as the mean. Arrows indicate direction of better results.

| Score function / model | EF<sub>1%</sub> ↑ | AUROC ↑ | BEDROC ($\alpha = 80.5$) ↑ |
| --- | --- | --- | --- |
| BIND | 24.14 | 0.952 | 0.698 |
| Zero-shot BIND (90% protein homology sequences removed) | 12.03 | 0.851 | 0.396 |
| SSM-DTA (pIC<sub>50</sub>) [14] | 10.19 | - | 0.336 |
| DeepRMSD [44] | 6.04 | - | - |
| OnionNet-SFCT with Vina [4] | 5.22 | 0.549 | - |
| Gnina [54] | 2.33 [4] | - | - |
| Vina [50] | 1.90 [4] | - | - |

\* best model's results as reported by respective authors

### 3.2 Protein language models can perform reverse screening

Reverse screening was performed on the benchmark as described by Luo *et al*., 2023 [17]. In summary, 90 selected ligands were scored against 12,195 human proteins representative of the human proteome. From the results, BIND is comparable to that of top models with rescoring such as OnionNet-SFCT with Glide-SP docking on AlphaFold2-predicted structures and PointSite or SiteMap for pocket determination; BIND being able to cumulatively rank the highest number of correct pairs for the top 2-1000 pairs (Fig. 4a-b). Furthermore, the reverse screening is expedient, taking only approximately eight hours on a Ryzen Threadripper Pro 5975WX CPU to run through all 1,097,550 protein-ligand combinations. The reverse docking performance of BIND was also evaluated on the Astex Diver dataset, of which in the top 1 scoring outperformed DOCK, Glide, and Autodock Vina, with comparable performance on the top 5 metrics (Fig. 4c).

**Figure 4.**
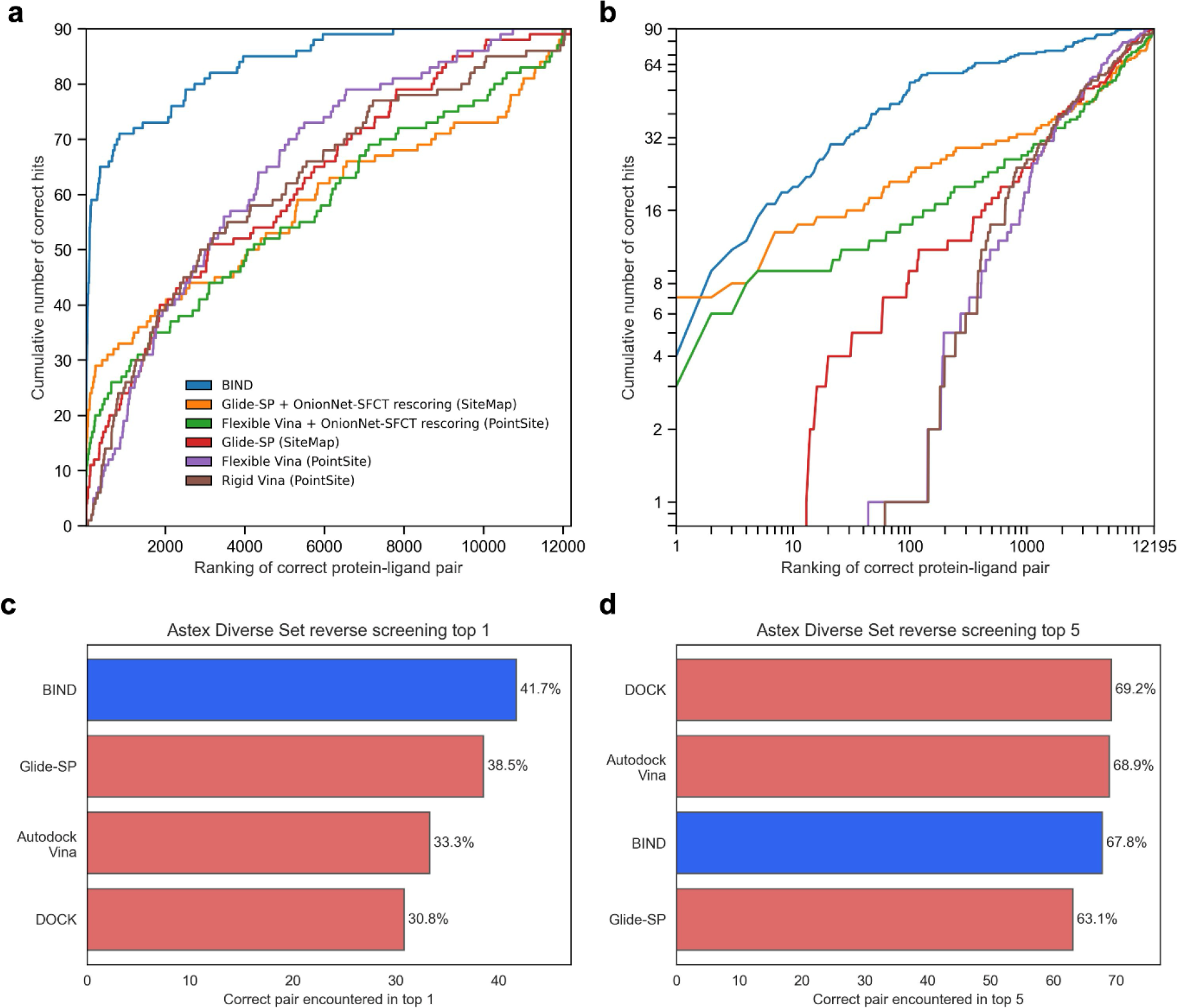
Protein language models can outperform standard reverse docking pipelines using structures from AlphaFold2. A total of 12,195 proteins and 90 ligands from Luo *et al*., 2023’s AlphaFold2 reverse docking benchmark were individually pairwise scored using BIND and the ligands ranked by classification score for each protein. (a, b) The cumulative ranking of BIND in standard and logarithmic plots respectively. The legend indicates the type of score function used, and the software used to determine the pocket locations in parentheses. (c, d) Ranking of 84 ligands against 85 proteins in the reverse docking benchmark on the Astex dataset, with other benchmark scores obtained from Luo *et al*., 2017.

### 3.3 Model can perform drug-target affinity prediction

The model is able to perform on the DTA prediction datasets DAVIS, despite not being specifically trained on this dataset but only BindingDB (Table 6). Furthermore, on the BindingDB test split, it achieved state-of-the-art RMSE in pK_d_ prediction specifically, but falls short in the other metrics of pK_i_, pIC_50_ and pEC_50_ (Table 7).

**Table 4.** Screening power results for DUD-E, with all scores represented as the mean.

| Score function / model | EF<sub>0.5%</sub> ↑ | EF<sub>1%</sub> ↑ | AUROC ↑ | BEDROC ($\alpha = 80.5$) ↑ |
| --- | --- | --- | --- | --- |
| BIND | 51.88 | 46.35 | 0.944 | 0.733 |
| RTMScore [5] | 42.47 | 35.10 | 0.831 | 0.558 |
| GenScore [42] | 44.02 | 33.96 | 0.828 | 0.546 |
| DyScore (target-aware split) [53] | - | 32.20 | 0.858 | 0.502 |
| PIGNet2 [43] | 36.80 | 31.20 | 0.850 | 0.515 |
| Zero-shot<br>BIND (90%<br>protein<br>homology<br>sequences<br>removed) | 30.52 | 26.39 | 0.826 | 0.430 |
| SSM-DTA<br>(pIC<sub>50</sub>) [14] | 33.59 | 24.78 | - | 0.397 |
| Gnina [54] | - | 20.40 [55] | 0.795 [55] | - |
| OnionNet-SFCT with<br>Vina [4] | - | 15.54 | 0.724 | - |
| DeepRMSD<br>with Vina* [44] | - | 14.80 | - | - |
| DyScore<br>(target-<br>unaware) [53] | - | 12.10 | 0.755 | 0.216 |
| Vina [50] | - | 8.82 [4] | 0.697 [4] | - |

\* best model's results as reported by respective authors

**Table 5.**
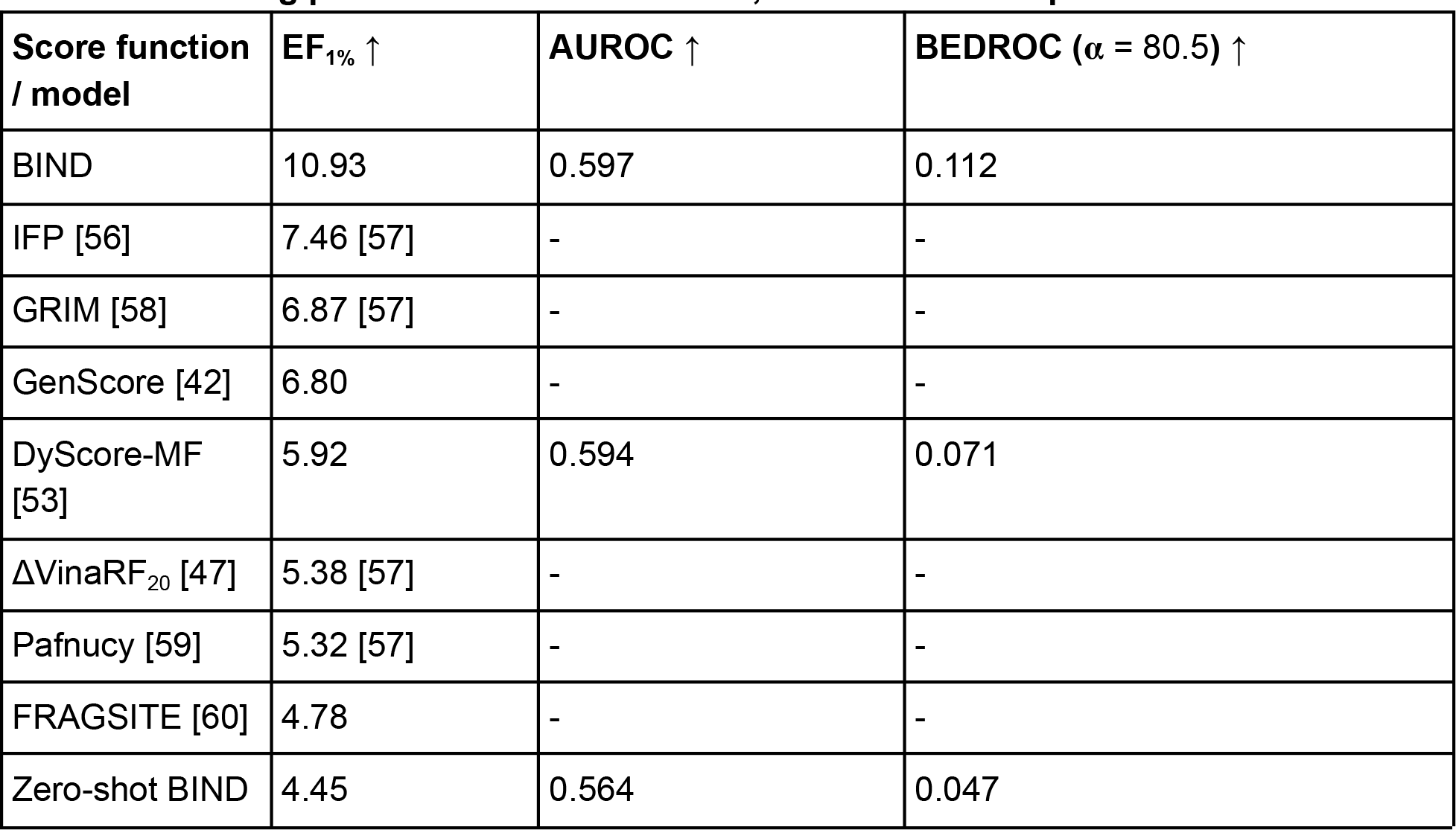
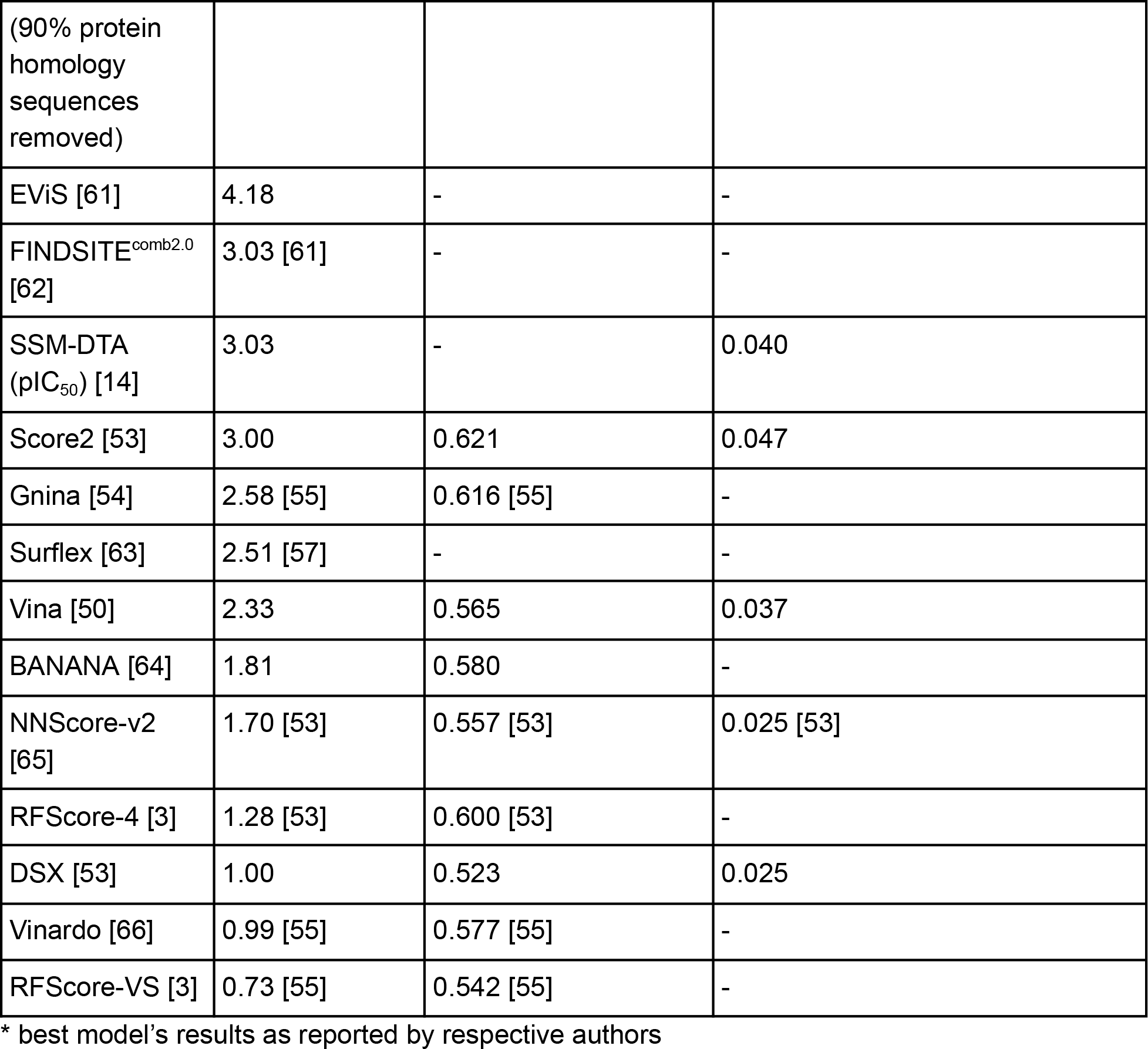
Screening power results for LIT-PCBA, with all scores represented as the mean.

**Table 6.** Drug-target affinity on benchmark datasets.

| **Dataset** | **DAVIS (pK<sub>d</sub>)** |  |
| --- | --- | --- |
| **Model** | **MSE ↓** | **CI ↑** |
| SSM-DTA [14] | 0.219 | 0.875 |
| DeepDTA [67] | 0.262 | 0.870 |
| ELECTRA-DTA [68] | 0.238 | 0.897 |
| SubMDTA [69] | 0.218 | 0.894 |
| DeepPurpose [70] | 0.242 | 0.881 |
| AttentionMGT-DTA [71] | 0.193 | 0.891 |
| AttentionDTA [72] | 0.195 | 0.888 |
| BIND (trained on BindingDB only) | 0.409 | 0.747 |

\* best model's results as reported by respective authors

**Table 7.** Drug-target affinity on BindingDB.

| **Metric** | **pK<sub>i</sub>** |  | **pK<sub>d</sub>** |  | **pIC<sub>50</sub>** |  | **pEC<sub>50</sub>** |  |
| --- | --- | --- | --- | --- | --- | --- | --- | --- |
| **Model** | **RMSE ↓** | **R ↑** | **RMSE ↓** | **R ↑** | **RMSE ↓** | **R ↑** | **RMSE ↓** | **R ↑** |
| DeepAffinity* [73] | 0.840 | 0.840 | - | - | 0.780 | 0.840 | - | - |
| SSM-DTA [14] | 0.792 | 0.863 | - | - | 0.712 | 0.878 | - | - |
| BACPI [74] | 0.800 | 0.860 | 1.080 | 0.730 | 0.740 | 0.860 | 0.780 | 0.850 |
| BIND | 0.965 | 0.798 | 0.935 | 0.798 | 0.924 | 0.808 | 0.891 | 0.822 |
| MONN [75] | - | - | - | - | 0.764 | 0.858 | - | - |

\* best model's results as reported by respective authors

### 3.4 Drug-target affinity prediction alone has lower screening power compared to classification

When ranked using pK_i_, pK_d_, pIC_50_ or pEC_50_, the enrichment factors across benchmark datasets dropped compared to classification alone (Fig. 5a - e). Furthermore, the classifier also has the strongest enriching effects in the AlphaFold2 reverse docking benchmark (Fig. 5f). Lastly, in spite of state-of-the-art DTA predictors being more accurate in predicting DTA compared to BIND (Table 6), BIND still achieves higher screening power on CASF-2016, DEKOIS 2.0, DUD-AD, DUD-E and LIT-PCBA using its pIC_50_ prediction compared to SSM-DTA’s pIC_50_ (Supplementary Fig. 2).

**Figure 5.**
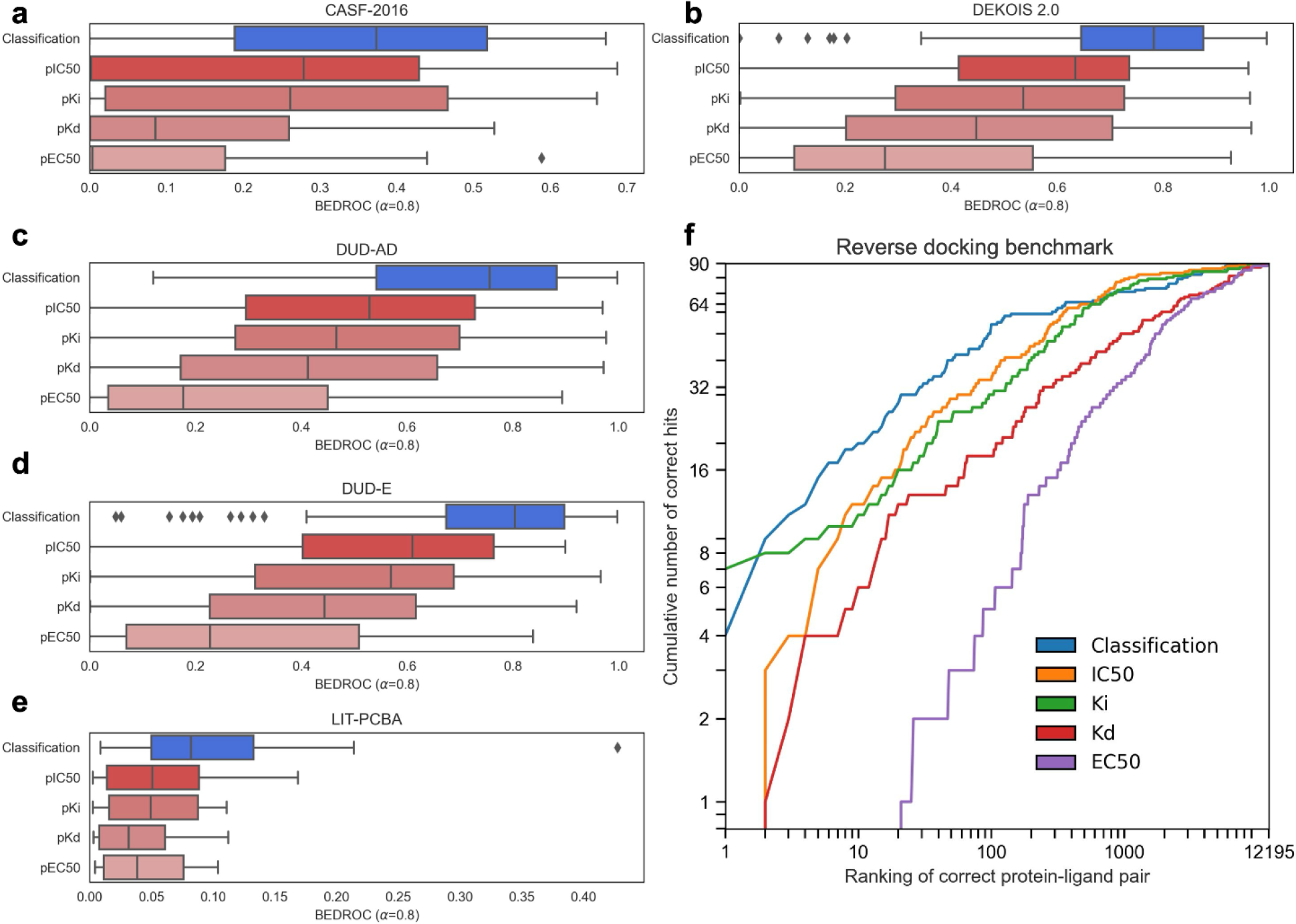
Predicted binding affinity has lower enriching power compared to the classifier in both forward and reverse screening. (a-e) Box-and-whisker plots of BEDROC values across the entire CASF-2016, DEKOIS 2.0, DUD-AD, DUD-E and LIT-PCBA datasets respectively, with boxes representing the interquartile range and the median demarcated in the box, whiskers showing the fence and diamonds showing outliers. (f) Cumulative logarithmic plot of ranking in reverse screening on Luo *et al*., 2023’s AlphaFold2 reverse docking benchmark dataset.

### 3.5 Attention weights can pinpoint residues important for binding despite the model not being trained for such

187 single-chain entries in the CASF-2016 core set were put into the model and the residues with the highest attention scores extracted, with attention scores maxed across all four cross-attention blocks. The model is able to identify the binding residues (Fig. 6), and hence determine potential binding pockets in a protein with unknown structure. At a 10Å cutoff, the model is able to identify approximately 30% of binding pocket residues. However, the attention weights are still patchy across the entire protein (Fig. 6b).

**Figure 6.**
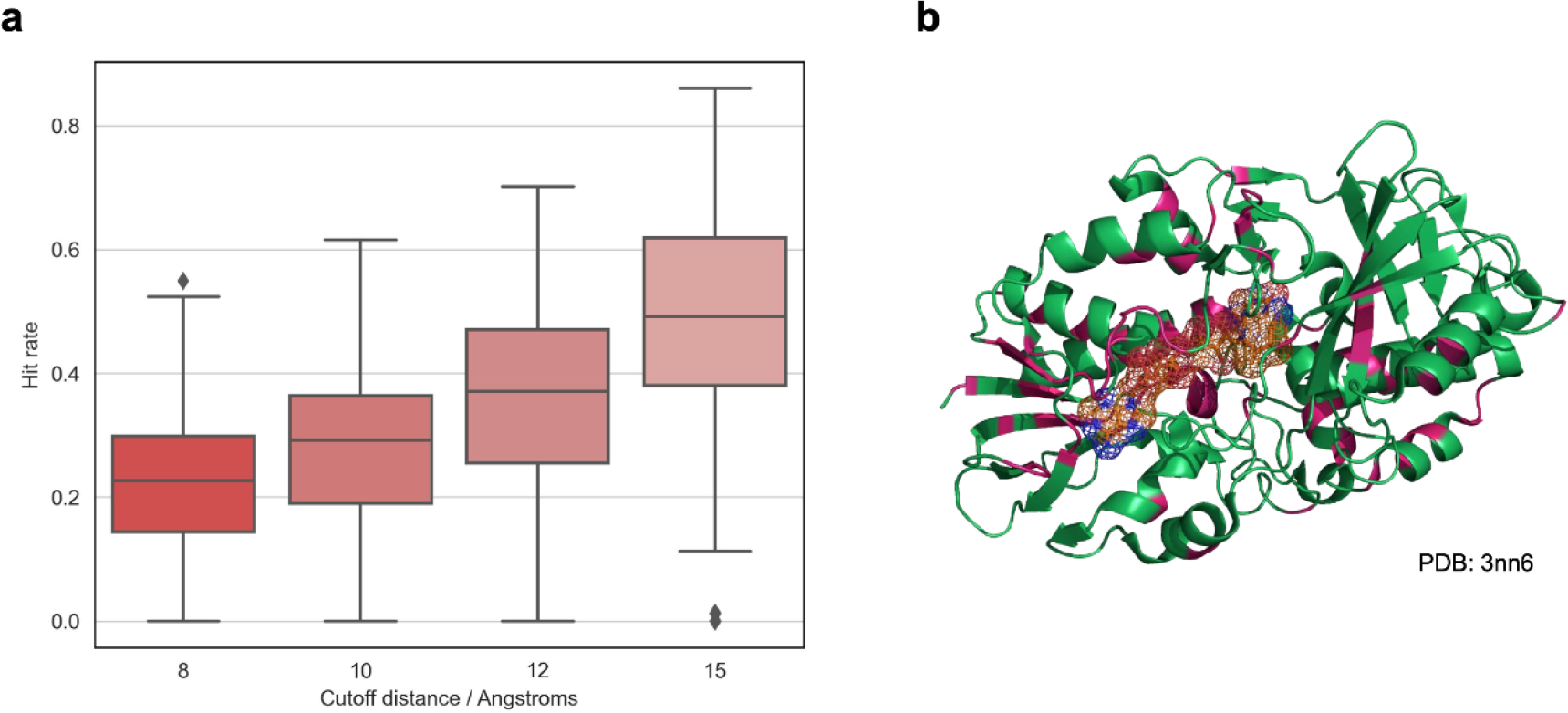
Pocket residues can be elucidated from the cross-attention scores of the model despite the model not being trained to do so. (a) 187 single-chained proteins from the CASF-2016 core set are inputted into BIND, with the attention scores averaged across all four cross-attention graph block layers. The proportion of binding pocket residues identified is shown at different cutoff distances. Boxes represent the interquartile range with the median demarcated in the box, whiskers showing the fence and diamonds showing outliers. (b) A visual example of the top 20% most attended to residues in 6-hydroxy-L-nicotine oxidase coloured in pink.

## 4. Discussion

The use of protein language models in predicting DTA is not new. Previous work has delivered very accurate DTA predictors using PLMs such as SSM-DTA [14]. However, there has been far less application of this to determine screening power - arguably the most important goal in CADD. Previous work by Tsubaki *et al*., 2019, used a one-dimensional n-gram convolutional neural network and cross-attentions to do similar work, achieving good discriminative power between real and decoy ligands when trained and evaluated on the DUD-E dataset using fivefold cross-validation [18]. This work builds upon that foundation and uses the pre-trained ESM-2 to transfer pre-learned knowledge about the protein language into the virtual screening pipeline.

Primarily, BIND serves as a proof of concept that even in the complete absence of protein structure or binding pocket information, it is possible to achieve screening power similar to that of and sometimes even exceed certain top SBDD models with PLMs by training the model to discriminate between real and decoy ligands selected from the same dataset. The results show that by using a model such as the proposed BIND, it may be possible to perform virtual screening for drug discovery, or to elucidate the identity of protein targets which bind to a specific ligand as per reverse docking. However, from the CASF-2016 results, it appears that BIND is not the best at ranking the very top and most potent binders as apparent in its poorer success rate in a much smaller set of ligands as seen in CASF-2016. From this, and with the comparable performance and occasionally state-of-the-art enrichment factor scores in the other larger datasets, it can be said that BIND has rather demonstrated its utility in enrichment of large set; and given the high speed at which BIND can screen compounds, it can be pragmatically deployed in initial screening stages to filter out a large number of non-binders (i.e. from millions to thousands), of which refinement can then be done with other score functions such as GenScore, PIGNet2 or OnionNet-SFCT to pull out only the top binders. The strong enriching effects of BIND in reverse screening also indicates that BIND could be better at finding a target protein from a large number of proteins given a ligand than performing docking on AlphaFold2 structures. This is especially useful as, in many reverse docking pipelines, the experimental structure of all proteins is typically not available, and reverse docking may be performed on non-binding protein conformations [19].

The authors posit from the results that BIND is able to achieve the reported screening power by borrowing the pre-learned context from PLMs such as ESM-2, which are able to understand the implicit structure and underlying language of proteins. This includes understanding biochemical properties, dynamic movement of proteins, and interdomain interactions.The performance of the zero-shot model also shows BIND has sufficient generalisation capability in screening unseen proteins, potentially obtaining this capability from its ESM-2 pre-training which allowed it to see a much more diverse set of protein sequences. This premise of stronger performance from transfer learning is supported by that of literature [20], and previous work such as ESMFold have demonstrated that structure can be inferred from sequence alone with the help of PLMs.

Therefore, it may not be far-fetched that the flexibility and even how a protein interacts with other molecules such as ligands be implicitly learnt through the masked language modelling and other self-supervised objectives of PLMs in pre-training. PLMs may be able to better model molecular interactions with disordered regions in proteins and even decipher cryptic pockets which evade the typical SBDD virtual screening pipeline due to their transient nature [21]. There remains much to be explored in this domain.

Furthermore, it is interesting that the BIND model’s DTA prediction has lower screening power and enriching effects compared to discriminating between true and false binders (i.e. the classifier in BIND) alone. It is highly plausible that this is due to internal biases of the training dataset - as all of the training protein-ligand pairs are true binders, the algorithm is misled into thinking that every protein-ligand pair that is given is always a binder and hence predicts a binders’ score regardless to prevent heavy loss penalties, as this is reflective of what it has seen and points raised by previous work. This result reinforces the indication that robust augmentation very likely goes hand-in-hand with screening power. The use of sole DTA models in screening may also not be appropriate in all cases, as DTA datasets, including BindingDB, which derives its affinity values from multiple sources, have been shown to be noisy across multiple assays [22] - this underscores that it may be more practical to predict a binary label as to whether or not a drug binds than a quantitative experimental affinity that is subject to many other confounding and unforeseen experimental variables, which could also explain BIND’s higher screening power compared to SSM-DTA although BIND achieves significantly worse R^2^ and MSE values in IC_50_ and other DTA predictions across the board.

The authors acknowledge several limitations of this work: (1) The BindingDB dataset is mainly obtained from ChEMBL and other sources, and contains overlapping information with the CASF-2016, DEKOIS 2.0, DUD-AD, DUD-E and LIT-PCBA evaluation datasets, in the process inflating the results for the forward screening for the non-zero-shot model. Although this is an issue, it is difficult to directly and objectively quantify the overlaps and individually eradicate the overlaps. Furthermore, by removing overlaps, a subjective defined stringency is required (i.e. Tanimoto similarity, protein sequence similarity), in the process getting rid of a large, diverse and representative chunk of protein-ligand binding data in *Homo sapiens* and other key protein families (which is the entire rationale of the benchmark datasets). This would prejudice BIND in evaluation as other score functions are typically trained with knowledge of that protein previously (i.e. from the PDBBind dataset [10], which has directly overlapping PDB structures with the evaluation dataset and are typically not removed during model training). By removing direct ligand-protein matches but not proteins during evaluation, the evaluation cannot be fairly performed with other score functions using the same dataset as the dataset would be modified and the enrichment factors cannot be directly compared. To mitigate this limitation and to assess the true screening power of BIND, the zero-shot model was attempted in which sequences containing > 90% homology on the proteins were removed from the training set, and the model still showed generalisation capability compared to contemporary score functions; (2) the explainability of BIND is limited - although the cross-attentions are generally able to pinpoint the important residues in a protein, they are far from perfect and are still largely uninterpretable in-line with post-hoc explainability analyses performed on transformer models; (3) much like previous work such as SSM-DTA, the training expense of this model is high. To compensate with limited resources, the authors used a large number of gradient accumulation steps, in the process dragging out the training time. The model may also further benefit in the DTA prediction criteria from a scaled-up training regime, which could not be achieved in this work due to limited computational resources; (4) No wet lab experimental validation was performed for this work - although BIND performs reasonably well on benchmarks as explored in this work, many factors ultimately affect the efficacy of a drug and experimental validation is needed.

Overall, this work proposes the use of PLMs as a potential alternative for SBDD virtual screening - showing that, even without structural information, is comparable to top-of-the-line SBDD models. The authors reinforce that this methodology be termed sequence-based drug discovery (SeBDD) [23]. SeBDD could be useful in work in which the protein structure cannot be accurately resolved, or when the binding pockets are unknown. In current screening pipelines, SeBDD can be used as an initial filter in a VS campaign, in which the top molecules from SeBDD are selected for standard SBDD docking and MD simulations. Due to the simplicity of SeBDD compared with SBDD (no need for docking, preparing the Gasteiger partial charges of ligands and targets, etc.), pipelines can also be greatly accelerated by its use.

Future work in this domain likely involves simulated annealing of molecules and integrating SeBDD with fragment-based drug discovery (FBDD), the use of synthons [24], and machine learning-based chemical spaces to explore very large amounts of chemical spaces quickly and accurately.

## 5. Methods

### 5.1 Model training, architecture, and training dataset

The BIND model was written and evaluated in Python 3.11 using PyTorch Geometric 2.4.0 with a PyTorch 2.2.0+cu121 backend and HuggingFace Transformers 4.37.2. The full model, along with its trained weights, is disclosed in the GitHub repository. Three models were trained with the one with the lowest validation loss selected and benchmarked in this study, with each model trained on a single Nvidia RTX A6000 for approximately 28 days. The whole model is described in Fig. 2. The implementation code for the cross-attention graph block is also in the same GitHub repository - in essence, each node in the graph is unwrapped and treated as an individual token, before a standard cross-attention similar to that of the original transformer decoder was used - the graph is then reconstructed afterwards. Five prediction heads are used in the final layer of the model, four of which are for regression objectives - to predict the pK_i_, pK_d_, pIC_50_, and pEC_50_ respectively. A classification objective is also added to predict which ligands are true and which are decoys. The logits of the classification head are multiplied by the natural exponential of a trainable temperature parameter, τ, initialised at 0.07 as per previous work [25]. After scaling, logits are clipped from -100 to 100 to ensure training stability. The loss functions used are Huber loss with a δ = 2.0 and binary cross entropy for regression and classification heads respectively. Training was performed using the AdamW optimiser with a learning rate of 1e-4 and a weight decay of 1e-3. A dropout of 0.1 was applied to the multi-headed attention layer, and a leaky rectified linear unit (ReLU) activation of ɑ = 0.05 is used unless otherwise specified. Learning rate was scheduled using cosine decay with no warm up. The 650 million parameter ESM-2 (facebook/esm2_t33_650M_UR50D) model’s weights were retrieved using the HuggingFace Transformers 4.37.1 library and are frozen during training. Latents from the ESM-2 layers 1, 11, 21, 31 are cross-attended to. Convolutions for the molecular graph are GATv2 [26] for all instances, and pooling used for the interaction network is Learnable Commutative Monoid (LCM) Aggregation [27]. Model training was done in automatic mixed precision and FlashAttention-2 [28] was used where applicable. The training dataset used is BindingDB [29], with a total of 2,469,626 (90%) protein-ligand pairs used for training, 54,902 (2%) for validation, and 224,738 (8%) for testing. A batch size of 1 was used during training together with 256 gradient accumulation steps, for a total of 100,000 full iterations (effectively around 10.37 epochs’ worth). To optimise for computation, all protein sequences exceeding 2,048 amino acids were removed. In total, 1,202,465 unique SMILES strings as part of the BindingDB dataset were used. Since there are multiple prediction heads, for entries in which one or more of the regression values were unavailable, the local loss was set to zero, meaning that there is no penalty for that term regardless of what the model predicts. For decoy ligands specifically, the loss for K_i_, K_d_, IC_50_ and EC_50_ terms are set to zero. All affinity values were normalised with -log_10_(x / 1e9), where x is in nM, consistent with all previous work [14]. For the reported training loss, the zeroised terms are disregarded in averaging. Approximately 50% of ligands used during training are decoys as part of a data-balanced regime.

### 5.2 Graph node and edge generation

The construction of the molecular graph is identical to that done in previous work [30], with the exception that the PySmiles library is now set to not reinterpret aromatic bonds due to a software bug. PySmiles 1.1.2 and NetworkX 3.2.1 were used to generate the graphs used by the model.

### 5.3 Decoy ligand determination

Decoys were determined at training by randomly selecting another ligand from the loaded SMILES, and programmatically checking if there is another entry in the BindingDB data that tags that SMILES to the same protein sequence.

### 5.4 Evaluation datasets

The DEKOIS 2.0 [31], DUD-E [32], DUD-AD [16], LIT-PCBA [33] and CASF-2016 [6] datasets were used to determine screening power. In all instances, ligands were converted into SMILES using OpenBabel 3.1.0. For all datasets, the UniProt sequences were obtained which corresponded to each RCSB PDB entry. In instances where multiple sequences relating to the target were present, the sequences were concatenated before inputting into the model - protein sequences that were unrelated to the test protein such as hirudin in thrombin and nuclear cofactors in the oestrogen receptors were removed. All protein sequences used in evaluation are available in Supplementary File 3. DTA evaluation was performed on the DAVIS [34] dataset obtained from TDCommons [35], and binding affinity prediction was also performed on a test set of BindingDB split prior to training. Unlike training, there is no cutoff on protein length used during evaluation/testing.

### 5.5 Dataset splitting and zero-shot protein performance evaluation

As there are overlaps between the BindingDB dataset and the screening power evaluation datasets, sequence-ligand pairs in BindingDB which contained > 90% overlap in the protein sequence compared to the UniProt sequences in the evaluating datasets were removed for the zero-shot model. The percentage alignment was calculated using Biopython 1.78’s pairwise2 align module, with the UniProt sequence used as the reference length in computing the final homology percentage. The zero-shot model was re-trained on this dataset with the exact same parameters as stated in 5.1.

### 5.6 Evaluation with SSM-DTA

The SSM-DTA [14] pIC_50_ model (BindingDB_IC50/checkpoint_best_20221021) was used on the same datasets described in 5.4, with the same protein sequence input and SMILES as per Supplementary File 3, after canonicalisation and tokenisation as provided by the SSM-DTA source code.

### 5.7 Metric calculation

BEDROC [36] and Enrichment Factors were calculated using RDKit 2023.09.5, whereas AUROC was calculated using the SciPy 1.11.0 library. Reported scores are averaged across all entries in the respective evaluation datasets (57 for CASF-2016, 81 proteins for DEKOIS 2.0, 102 for DUD-AD and DUD-E, 15 for LIT-PCBA). All decoys and ligands are used in benchmarking as per the original datasets.

### 5.8 Reverse screening

Reverse screening was performed as per Luo *et al.*, 2023 [17]. In essence, 90 ligands were selected and reverse screened on a trimmed down human proteome of 12,195 proteins from UniProt using BIND. For the reverse docking performed using SBDD models done by Luo *et al*., 2023, structures obtained from the AlphaFold Protein Structure Database [37] were used. PointSite [38] or SiteMap [39] were used to determine potential binding pockets. Reverse screening was also done on the Astex Diverse Set using the same procedure as with other standard docking tools described by Hartshorn *et al.*, 2007 [40]. Summarily, the 84 ligands found in the dataset were docked against the 85 proteins to identify the correct protein target for the ligand and ranked, with the percentage of correct pairs identified in the top 1 and top 5 rankings as per literature [41].

### 5.9 Attention mapping and scoring

Attention weights were extracted per layer and averaged over all heads as per the MultiHeadedAttention class in PyTorch. Subsequently, they are averaged across the four cross-attention graph blocks in BIND, and the maximum weight per residue selected across all atoms, creating a single attention weight score per residue between 0-1.

### 5.10 Attention scores binding pocket overlap determination

The CASF-2016 core set was filtered for single chains. Of which, 187 out of 285 proteins fulfilled aforementioned criteria. Open-source PyMOL 2.5.0 was then used to determine all residues within 8Å, 10Å, 12Å and 15Å of the native ligand as provided by CASF-2016 respectively. Residues are counted so long as any part of the amino acid (including its sidechains) are within the cutoff distance to any of the ligand atoms using the “X within Y of Z” command in the PyMOL API. Attention scores from BIND were determined as previously described, and the top number of residues corresponding to the total number of residues selected with respect to the docked ligand (through PyMOL) were extracted and overlaps counted. In essence, this metric measures the proportion of most attended residues to the actual binding pocket. Generally, the larger the cutoff value, the easier it is to get a higher metric score.

## Acknowledgements

The authors thank Saxena Shikhar for the fruitful discussions as part of this work. We further would like to express our appreciation to Qing Luo and Jingjing Guo of Macao Polytechnic University for supplying their full reverse docking data from their previous work.

## 6. Code Availability

The code is available at: https://github.com/Chokyotager/BIND.

## 8 Funding

This work is supported by the Singapore Ministry of Education (MOE) Tier 1 grant RG97/22. The computational work for this article was partially performed on resources of the National Supercomputing Centre, Singapore (https://www.nscc.sg) and the HADLEY supercomputer by Singapore Centre for Environmental Life Sciences Engineering (SCELSE). SCELSE is funded by Singapore’s National Research Foundation, the Ministry of Education, Nanyang Technological University (NTU), and the National University of Singapore (NUS), and is hosted by NTU in partnership with NUS.

---

**Supplementary Figure 1.**
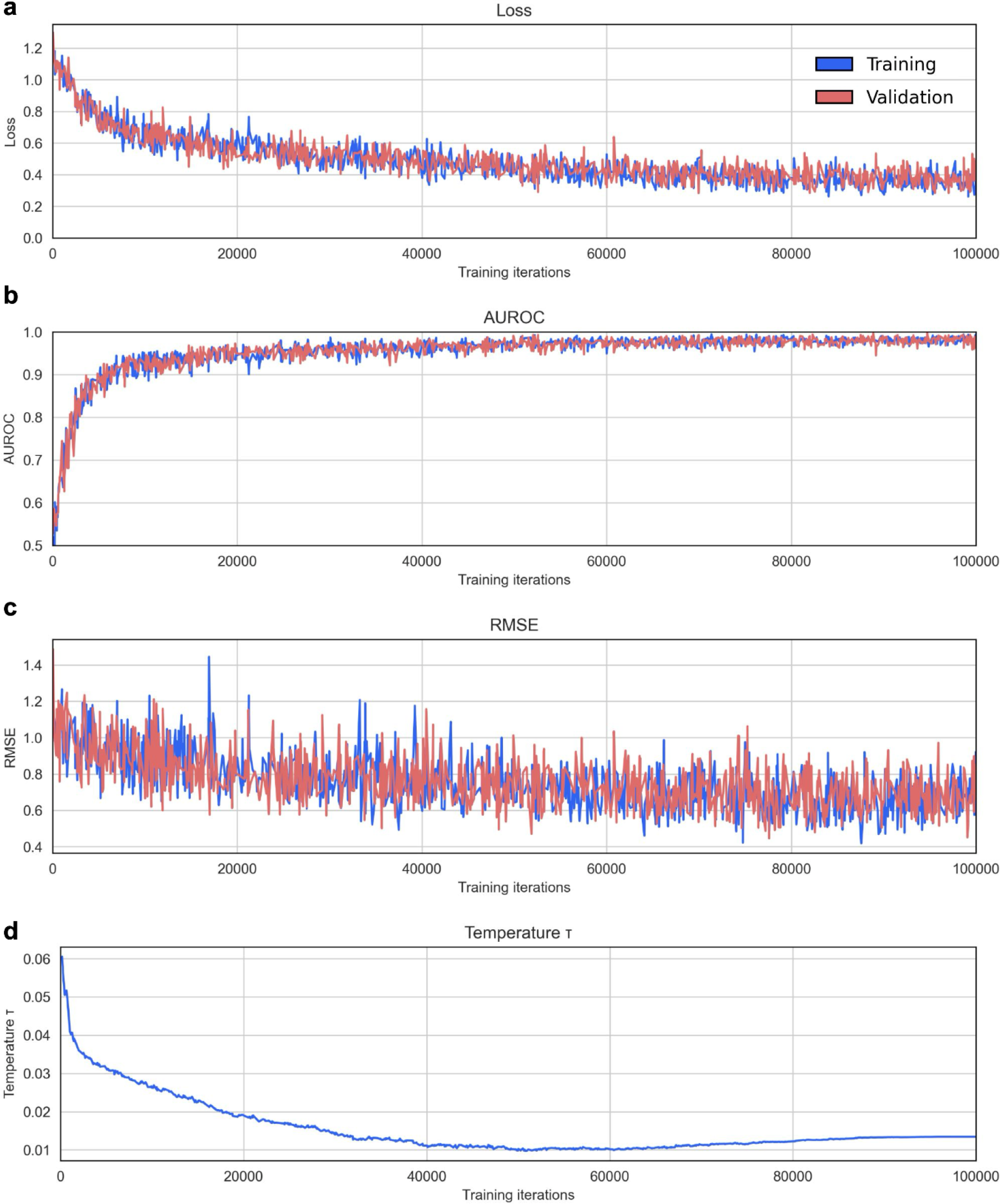
Training and evaluation metrics for the model. (a) Loss curve with a combined loss of Huber for the DTA regression and binary cross entropy for decoy/true binder classification; (b) AUROC train and validation curve; (c) DTA RMSE train and validation curve; (d) temperature parameter throughout training.

**Supplementary Figure 2.**
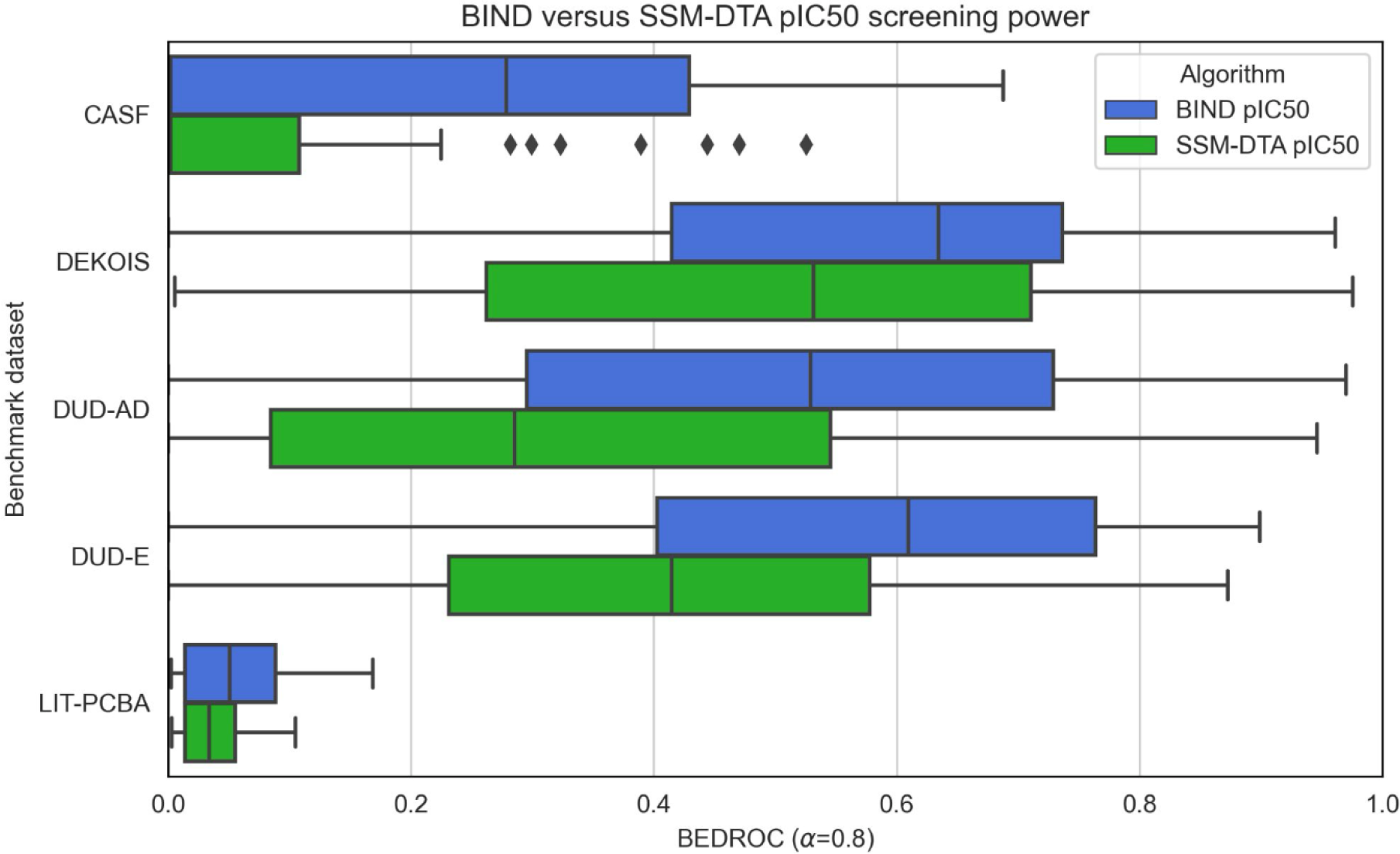
BIND still achieves higher screening power than SSM-DTA on its pIC_50_ prediction despite SSM-DTA’s being more accurate when measured via the RMSE metric. The BEDROC values across the CASF-2016, DEKOIS, DUD-AD, DUD-E and LIT-PCBA datasets are compared and calculated by ranking on the pIC_50_ predictions of both BIND and SSM-DTA. Boxes represent the interquartile range with the median demarcated in the box, whiskers showing the fence and diamonds showing outliers

**S1 Model experiments.** In developing the model, different model configurations were trialled. The final, most successful version is described under the methods section of this work. Notably, flipping the order of ESM-2 latents input (i.e. cross-attending to downstream parts of the ESM-2 transformer before upstream) into the model resulted in worse results. Other experiments included having the ESM-2 embeddings cross-attend to the molecular graph instead, and then mean pooling over the tokens before a small feedforward layer. These resulted in similar RMSE predictions to having the molecular graph query the ESM-2 embeddings but drastically increased the computational overhead. When training with lower gradient accumulation steps of 128, certain instances of the model failed to converge. Lastly, an implicit hydrogen model was attempted, however these failed to perform as well as the explicit hydrogen models. The authors hypothesise that the addition of hydrogen atoms resulted in lesser compression of information in the form of latent space in the model, and hence culminated in better results overall.

**Supplementary File 1. All evaluation metrics across individual targets in different evaluation datasets using BIND.**

**Supplementary File 2. All evaluation metrics across individual targets in different evaluation datasets using zero-shot BIND.**

**Supplementary File 3. Protein sequences used in evaluation on CASF-2016, DEKOIS 2.0, DUD-AD, DUD-E, and LIT-PCBA.**

## Notes

### Competing Interest Statement

The authors have declared no competing interest.

### Summary of Updates

New results regarding zero-shot model; limitations in discussion updated; new section in methods regarding the zero-shot model; Figure 3 updated; One additional supplementary file.

